# Microstate-specific functional connectivity in alcohol use disorder using resting-state EEG

**DOI:** 10.64898/2026.01.15.699621

**Authors:** Hanjin Park, So Young Yoo, Hyunho Lee, Minkyung Park, Jung-Seok Choi, Dongil Chung

## Abstract

Alcohol Use Disorder (AUD) is often described as a “disconnection syndrome”, reflecting neural dysfunction associated with chronic alcohol use. However, previous studies on functional connectivity of resting-state EEG (rsEEG) in AUD have reported inconsistent findings—some showing hyperconnectivity, others hypoconnectivity or no significant differences—possibly due to overlooking the brain’s dynamic nature even at its resting state. Here, we examined rsEEG in 26 young adults with AUD and 35 healthy controls (HC), using microstate analysis to characterize transient brain states and to estimate functional connectivity within each state. We showed that microstate map topologies were comparable between the two groups. However, the temporal dynamics in the AUD group were biased toward microstates C and D, characterized by anterior-to-posterior configurations, within which between-channel functional connectivity differed from that of HC. Particularly within microstate C, the AUD group showed significantly reduced functional connectivity compared to the HC group in the majority of channel pairs. Moreover, compared with conventional functional connectivity estimates, microstate-specific functional connectivity yielded superior classification performance in distinguishing individuals with AUD from HC. Subsequent computation of graph-theoretical measures revealed that individuals with AUD showed less stable hub-like properties and reduced small-worldness in microstate C, indicating diminished network efficiency. Our findings suggest that transient EEG properties, exemplified by microstate-specific functional connectivity, provide informative neural markers for characterizing the impact of alcohol use on the brain.

**Highlights:**

- Resting-state EEG microstates reveal altered brain dynamics in alcohol use disorder
- Microstate C exhibits widespread reductions in functional connectivity in AUD
- Microstate-specific connectivity outperforms conventional measures in AUD classification
- Graph analysis reveals reduced network efficiency and unstable hubs in AUD

## Introduction

Alcohol Use Disorder (AUD) is a chronic condition characterized by neural dysfunction, leading to alterations in brain structure and function (Andrews-Hanna et al., 2007; Bottino et al., 2025; Chanraud et al., 2011; Dupuy and Chanraud, 2016). Chronic alcohol use, associated with impaired self-control and impulsivity behavior, is often described as “disconnection syndrome“—a condition characterized by disrupted neural connectivity (Dupuy and Chanraud, 2016; Elton et al., 2021; Shokri-Kojori et al., 2017; Vergara et al., 2017). In line with such notion, previous studies analyzed synchronization and phase locking value (PLV) of resting-state electroencephalography (EEG) in AUD and showed that their functional connectivity is impaired (de Bruin et al., 2004; Herrera-Díaz et al., 2016; Wang et al., 2025). Other neuroimaging studies corroborated these reports and suggested that disruptions in functional connectivity within the default mode network (DMN) may underlie the behavioral impairments observed in individuals with AUD (Andrews-Hanna et al., 2007; Bottino et al., 2025; Chanraud et al., 2007; Chanraud et al., 2011). Despite a large body of accumulated data and evidence, there have also been studies in AUD reporting findings that contradict these results, reporting either hyperconnectivity (Winterer et al., 2003) or no statistical differences in functional connectivity compared to healthy controls (HC) (de Bruin et al., 2004; Park et al., 2017; Sion et al., 2020). Such inconsistencies may be explained by the well-known large individual differences observed among individuals with AUD (Andrews-Hanna et al., 2007; Bottino et al., 2025; Chanraud et al., 2011; Dupuy and Chanraud, 2016), although alternative possibilities have not yet been explored.

Many classical EEG approaches, including those used in various AUD studies, assume stationarity in EEG signals, meaning that statistical properties such as the mean and variance remain constant throughout the analysis window. In contrast to this statistical assumption, another line of research treats the EEG signal as a dynamical system that transitions among multiple transient brain states (Betzel et al., 2012; Khanna et al., 2015; Tarailis et al., 2024). Analyses of the state occurrences and transition properties have been linked to various neurodegenerative (Nishida et al., 2013; Stevens and Kircher, 1998) and neuropsychiatric disorders (Kikuchi et al., 2011; Kim et al., 2024; Strik et al., 1995). Given that relationships among signals with dynamic properties may be over- or underestimated depending on their respective states (Stam, 2005), previously reported inconsistencies in functional connectivity findings in AUD may stem from overlooking such dynamic state changes.

Here, we propose that integrating microstate analysis, which classifies transient brain states with distinct scalp topographies, with the computation of functional connectivity within each state would provide a more stable tool for examining potential neural alterations in individuals with AUD. To test this possibility, we analyzed resting-state EEGs (rsEEGs) recorded from 26 young adults with AUD and 35 healthy controls (HC), identified latent microstates to account for temporal dynamics in the signals, and computed functional connectivity associated with each state. Subsequently, both the temporal properties of microstates and the graph-theoretical network properties were compared between the two groups, and their contributions to classification accuracy were tested against those of conventional functional connectivity approaches.

## Methods

### Participants

Twenty-six young adults diagnosed with AUD (male/female = 19/7, age = 27.62 ± 5.39 years) and 35 healthy controls (male/female = 25/10, age = 24.94 ± 2.90 years) were recruited for the current study. Note that the age range of participants in the current study is 19-35, which contrasts with most previous AUD research that predominantly focuses on middle-aged populations (between 40 and 50) (Bottino et al., 2025). This age range was selected to reflect the earlier age at which individuals have been initiating alcohol consumption, as documented over the past two decades (Hingson et al., 2006), and to minimize potential confounding effects of comorbidity that become more common with aging (Sullivan and Pfefferbaum, 2019). AUD patients were recruited from the outpatient clinic at Seoul Metropolitan Government Seoul National University (SMG-SNU) Boramae Medical Center, while HCs were recruited from local communities and universities through public announcements. AUD was diagnosed by a clinically experienced psychiatrist according to DSM-5 criteria, and the individual patient’s severity was assessed using the Alcohol Use Disorders Identification Test (AUDIT) (Kim et al., 2008; Saunders et al., 1993). HCs consumed no more than 14 standard drinks per week and no more than four standard drinks on any single occasion. HC participants had no prior history of AUD or psychiatric disorders. None of the participants had a history of intellectual disability, psychotic, neurological, or seizure disorders. Furthermore, all participants were right-handed, medication-naïve, and had an intelligence quotient (IQ) above 70. The study protocol was approved by the Institutional Review Board of the Seoul Metropolitan Government Seoul National University (SMG-SNU) Boramae Medical Center, Seoul, Republic of Korea (IRB number: 16-2014-139), and all procedures were conducted in accordance with the Declaration of Helsinki. All participants provided written informed consent before participation.

### Clinical measures and psychological assessments

Demographic information, including age, sex, and years of education, was collected from all participants. The AUDIT (Kim et al., 2008; Saunders et al., 1993), a 10-item questionnaire that assesses hazardous drinking behavior, was administered to evaluate the severity of alcohol consumption. Each item was rated on a four-point Likert scale (0 = never, 4 = very frequent), and total AUDIT scores range from 0 to 40. Given the strong association between AUD and altered psychosocial status, we administered additional psychological questionnaires and used their scores as measures of individual differences. Specifically, individuals’ dispositional sensitivity to rewards and punishments was assessed using the Behavioral Activation Scale (BAS) and Behavioral Inhibition Scale (BIS) (Carver and White, 1994), which consist of 13 and 7 items rated on a four-point Likert scale, respectively. In addition, individuals’ stress levels and mental well-being were measured using the Psychosocial Well-being Index (PWI), a 45-item self-report questionnaire designed to evaluate physical and psychological well-being over the past several weeks (Kim, 1999). The total BAS score ranges from 13 to 52, the total BIS score ranges from 7 to 28, and the PWI score ranges from 0 to 135.

### EEG acquisition

The EEG data were digitized at a sampling frequency of 1000 Hz and amplified using SynAmps 2 (Compumedics, Abbotsford, Victoria, Australia) and the Neuroscan system (Scan 4.5, Compumedics). EEG signals were recorded with a 64-channel Quick-Cap (Neuroscan, Compumedics), which follows the modified 10–20 system for electrode placement. The recording took place in a sound-shielded room, and participants were instructed to keep their eyes closed for 5 minutes. The ground electrode was positioned between FPz and Fz, and a bipolar reference electrode was placed at the mastoid. Vertical and horizontal electrooculograms (EOGs) were recorded using electrodes positioned above and below the left eye to monitor ocular movements. The impedance of all electrodes was maintained below 5 kΩ throughout the recording. The collected EEG data were bandpass filtered between 0.1–100 Hz.

### EEG preprocessing

Raw EEG data were preprocessed using MNE-Python (Gramfort et al., 2013). For analysis, 19 channels (FP1, FP2, F3, F7, FZ, F4, F8, C3, CZ, C4, P3, PZ, P4, O1, O2, T7, T8, P7, P8) were selected from the 64-channel recordings to improve computational efficiency. To minimize potential edge artifacts, the first and last 30 seconds of the 5-minute recordings were excluded.

Preprocessing steps included bandpass filtering (0.5–51 Hz), re-referencing to the common average, segmenting the data into 1-second epochs, and rejecting epochs with amplitudes exceeding 100 μV. Independent component analysis (ICA) was performed to remove artifacts; however, upon visual inspection, no clear noise components were identified. Finally, an alpha-band (8–13 Hz) bandpass filter was applied, as this frequency range exhibited the highest global explained variance (GEV) in the subsequent microstate analysis (see below).

### Microstate analysis

To estimate microstates and their transitions, we conducted both individual- and group-level analyses. At the individual level, we first computed the global field potential (GFP) from the preprocessed EEG data and identified its local maxima. A modified k-means clustering algorithm (*k* = 4) was then applied to these GFP peaks, with 50 runs of cross-validation. Note that the number of microstate classes was fixed at four, consistent with the most commonly reported microstate patterns in the literature (Khanna et al., 2015; Tarailis et al., 2024). This procedure yielded four representative microstate clusters for each participant. At the group level, the individual-level microstate clusters were used as input to a modified k-means clustering algorithm to derive group-level microstate maps for the HC and AUD groups separately. With visual inspection, we confirmed that the microstate maps estimated for both groups corresponded well to the canonical microstates A, B, C, and D, i.e., the four prominent microstate classes typically reported in healthy individuals (Khanna et al., 2015; Tarailis et al., 2024). Subsequently, we computed cosine similarity to examine topological similarities between the two groups.

The template microstate maps (A-D) were then back-fitted to each participant’s preprocessed EEG data using a winner-takes-all approach with 35 msec temporal smoothing (Poulsen et al., 2018). This procedure yielded individual-level microstate sequences representing the latent dynamic EEG states for each subject. To characterize these sequences, we calculated the GEV (range = 0-1), which quantifies how well each of the four template maps accounts for the original EEG data based on their spatial correlation (Khanna et al., 2015). In addition, we computed two temporal parameters: time coverage, defined as the proportion of the total recording time occupied by each microstate, and mean duration, defined as the average continuous time each microstate remained active (Khanna et al., 2015).

### Microstate-specific functional connectivity

To assess microstate-specific functional connectivity, each individual’s EEG were segmented using a non-overlapping sliding time window. Specifically, we used a 230 msec time window to ensure coverage of at least three cycles of the fastest oscillation within the signal-of-interest (here, 13 Hz), allowing for reliable functional connectivity estimation (Cohen, 2014). Among the segmented time windows, we included only those in which a given microstate persisted for at least 210 msec, accounting for the 35-msec temporal smoothing applied during back-fitting, to ensure stable microstate-specific functional connectivity estimation.

Subsequently, the Hilbert transform was applied to EEG data from each selected window to obtain the instantaneous amplitude and phase. Coherence (COH) between channels *x* and *y* was defined as:

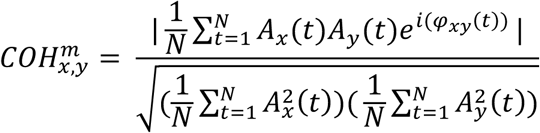

where *m* denotes a specific microstate, *A_x_*(*t*) is the instantaneous amplitude of channel *x* at time *t*, *φ_xy_*(*t*) is the instantaneous phase difference between channels x and *y* at time *t*, and *N* is the total number of time points. Coherence was computed between all possible channel pairs. To eliminate potential spurious coherence, coherence values lower than the median of all possible channel-pair coherence values were set to zero (Rubinov and Sporns, 2010). For each microstate-specific functional connectivity, coherence values were averaged across all selected windows belonging to that microstate.

As a comparison, conventional functional connectivity was also estimated from the EEG data without distinguishing microstates. For each participant, the EEG data were segmented into 1-second epochs, and coherence between all channel pairs was computed using the same procedure.

To evaluate the impact of distinguishing latent states in EEG functional connectivity estimation, we conducted classification analyses between the HC and AUD groups using a support vector machine (SVM) with a second-degree polynomial kernel. Specifically, the upper-triangular elements of the functional connectivity matrices were vectorized and z-scored to use as input features, and the participants’ group labels (HC or AUD) served as the output classes. Classification accuracy for the microstate-specific functional connectivity was evaluated using 5-fold cross-validation and was then compared with the classification accuracy obtained from the conventional functional connectivity.

For visualization, principal component analysis (PCA) was applied to the feature vectors, and the first two principal component scores were used to project the data into a two-dimensional space. In addition, to assess the distinctiveness of functional connectivity patterns across four microstates, we applied PCA-clustering to microstate-specific functional connectivity features within each group.

### Graph-theoretical network analyses

Previous studies suggest that graph-theoretical network analysis successfully captures alterations in neural mechanisms associated with brain dysfunction (Bassett and Sporns, 2017; Stam, 2014; Van den Heuvel and Sporns, 2013). To assess differences in functional network properties between the HC and AUD groups, the microstate-specific functional connectivity among EEG channels was modeled as a network, in which the channels served as nodes and the coherence between channel pairs served as edge weights. Prior to graph-theoretical analyses, weak connections—defined as edges with weights below the median of the connectivity matrix—were excluded to mitigate the impact of noise and low-reliability edges (Rubinov and Sporns, 2010).

Higher coherence values indicate stronger connections between nodes, which conceptually correspond to shorter distances in the network, reflecting more efficient information transfer between nodes (Rubinov and Sporns, 2010). Since Dijkstra’s algorithm requires the adjacency matrix to encode distances, the functional connectivity matrix was inverted before computing the shortest paths (Rubinov and Sporns, 2010). The shortest path between two nodes was then defined as the route with the minimal sum of edge weights.

To assess the efficiency of information communication within each network, we also identified hub nodes, which play a central role in integrating information across distant regions (‘integration’) and organizing functionally specialized sub-networks (‘segregation’) (Rubinov and Sporns, 2010; Van den Heuvel and Sporns, 2013). Specifically, we first identified candidate hub nodes as those whose strength (STR)—defined as the sum of a node’s weighted degrees—exceeded the mean plus one STD. We then examined four additional graph-theoretical measures: eigenvector centrality (EC), which evaluates a node’s influence by considering both its direct connections and the centrality of its neighbors; clustering coefficient (CC), which captures the local interconnectedness among a node’s neighbors; betweenness centrality (BC), which quantifies how frequently a node lies on the shortest paths between other nodes; and closeness centrality (C), which assesses a node’s efficiency in reaching all other nodes in the network (Sporns et al., 2007). Candidate nodes exceeding the threshold (mean plus one STD for EC, BC, and C; mean minus one STD for CC) in at least three out of the four graph-theoretical measures were identified as hubs.

- EC is defined as the component of the eigenvector corresponding to the largest eigenvalue of the adjacency matrix (Ruhnau, 2000).
- CC quantifies the fraction of triangles formed around a given node and represents the proportion of its neighbors that are also connected to each other (Rubinov and Sporns, 2010). Lower CC values suggest that the node functions as a bridge between distinct network regions.
- BC of a node *i* is defined as the fraction of all shortest paths between pairs of nodes that pass through the node *i*:

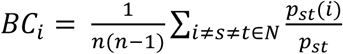 where *N* denotes the set of all nodes in the network, *n* is the total number of nodes, *p_st_* is the total number of the shortest paths between a node *s* and *t*, and *p_st_(i)* is the total number of the shortest paths between nodes *s* and *t* that pass through the node *i* (Rubinov and Sporns, 2010).
- C of a node *i* is computed as the reciprocal of the mean shortest path length from the node *i* to all other nodes in the network:

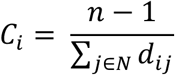

where *d_ij_* denotes the shortest path length between nodes *i* and *j*, and higher values indicate greater accessibility and more efficient information transfer across the network (Rubinov and Sporns, 2010).

In addition, we estimated small-worldness (SW), a well-established measure of efficient communication in complex networks (Rubinov and Sporns, 2010). SW was defined as:

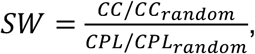

where *CC* and *CPL* denote the clustering coefficient and characteristic path length of the empirical network, respectively, and the *CC_random_* and *CPL_random_* represent the mean clustering coefficient and characteristic path length of the corresponding random networks. The CPL was calculated as:

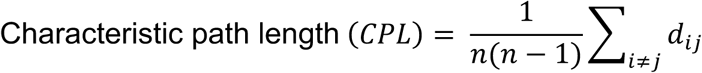

where *n* is the number of nodes in the network and *d_ij_* is the shortest path length between nodes *i* and *j*. To estimate the properties of random networks, we generated 500 degree-matched random networks for each empirical network and computed their mean *CC* and *CPL* values for comparison (Sporns et al., 2007).

All graph-theoretical measures were computed using the Brain Connectivity Toolbox (Rubinov and Sporns, 2010) in MATLAB.

### Statistical analyses

To investigate temporal dynamics across microstates within each group, we conducted repeated-measures analyses of variance (rmANOVAs) separately for each group and for each microstate parameter (i.e., GEV, time coverage, and mean duration), with microstate (A, B, C, and D) as a within-subject factor. Greenhouse-Geisser, Huynh-Feldt, and lower-bound corrections were applied to account for potential violations of the sphericity assumption. Post-hoc pairwise comparisons were conducted using Tukey’s Honestly Significant Difference (HSD) test to identify significant differences between microstates for each parameter within each group. Two-sample t-tests were conducted to examine group differences between the AUD and HC groups. Note that all between-group comparisons were controlled for individuals’ age. Multiple comparisons were corrected using the Benjamini-Hochberg false discovery rate (FDR) procedure (Benjamini and Hochberg, 1995), and a corrected p-value of less than 0.05 was considered statistically significant. Pearson’s correlation coefficients were computed to investigate the relationship between psychological variables and microstate-specific functional connectivity values. All statistical analysis was done using MATLAB R2021b (Mathworks).

## Results

### Demographics

The AUD and HC groups showed a comparable sex ratio (χ^2^(1) = 0.021, *P_FDR_* = 0.885), but differed significantly in age (t(59) = 2.12, *P_FDR_* = 0.019) and education (t(59) = -2.94, *P_FDR_* = 0.006) (**Table 1**). To control for age difference between the two groups, partial correlation analyses were subsequently conducted when examining group differences in microstate-specific functional connectivity measures and psychological variables. Because the EEG recordings in the current study did not involve specific cognitive tasks (i.e., rsEEG), we assumed that participants’ education level would not affect the measures-of-interest (e.g., functional connectivity) and therefore did not include it as a covariate.

**Table 1.** Demographic and psychological characteristics of the participants.

| | AUD (N = 26) | HC (N = 35) | Statistics | $P_{FDR}$-value |
| --- | --- | --- | --- | --- |
| Male/female<br>[% female] | 19/7 [26.9] | 25/10 [28.5] | $\chi^2(1) = 0.021$ | 0.885 |
| Age | 27.62 ± 5.39 | 24.94 ± 2.90 | t(59) = 2.12 | 0.019* |
| Education | 13.62 ± 1.44 | 14.94 ± 1.90 | t(59) = -2.94 | 0.006** |
| AUDIT | 27.77 ± 7.05 | 4.46 ± 3.01 | t(59) = 17.24 | 0.000*** |
| BIS | 20.65 ± 3.76 | 17.60 ± 3.84 | t(59) = 3.05 | 0.005*** |
| BAS | 34.69 ± 7.59 | 32.20 ± 6.28 | t(59) = 1.38 | 0.173 |
| PWI | 64.81 ± 26.03 | 29.89 ± 16.57 | t(59) = 6.28 | 0.000*** |

Values are presented as mean ± SD. AUD, Alcohol Use Disorder; AUDIT, Alcohol Use Disorder Identification; BIS, Behavioral Inhibition Scale; BAS, Behavioral Activation Scale; HC, Healthy Control; PWI, Psychosocial Well-being Index. \* $P_{FDR} < 0.05$ , \*\* $P_{FDR} < 0.01$ , \*\*\* $P_{FDR} < 0.001$ .

Consistent with the planned inclusion criteria, the AUD group showed significantly higher AUDIT scores than the HC group (t(59) = 17.24, *P_FDR_* < 0.001). Individuals with AUD are known to exhibit altered psychological characteristics, including heightened sensitivity to reward and punishment, and elevated stress levels (Andrews-Hanna et al., 2007; Bottino et al., 2025; Chanraud et al., 2011; Dupuy and Chanraud, 2016). In line with this literature, AUD showed significantly greater sensitivity to punishment (i.e., BIS: t(59) = 3.05, *P_FDR_* = 0.005) and higher PWI scores (t(59) = 6.28, *P_FDR_* < 0.001) compared to HC, while showing comparable sensitivity to reward (i.e., BAS scores) (t(59) = 1.38, *P_FDR_* = 0.173). Given that these measures capture psychological characteristics of AUD, we used them to examine individual variability within the AUD group.

### Individuals with AUD showed altered temporal properties across distinct microstates

For each group, four distinct microstate map topologies (microstates A, B, C, and D) were identified (**Fig. 1**). Cosine similarity analyses for each microstate indicated that the topologies were highly comparable between the two groups (A: 0.99, B: 0.97, C: 0.98, D: 0.95).

**Figure 1.**
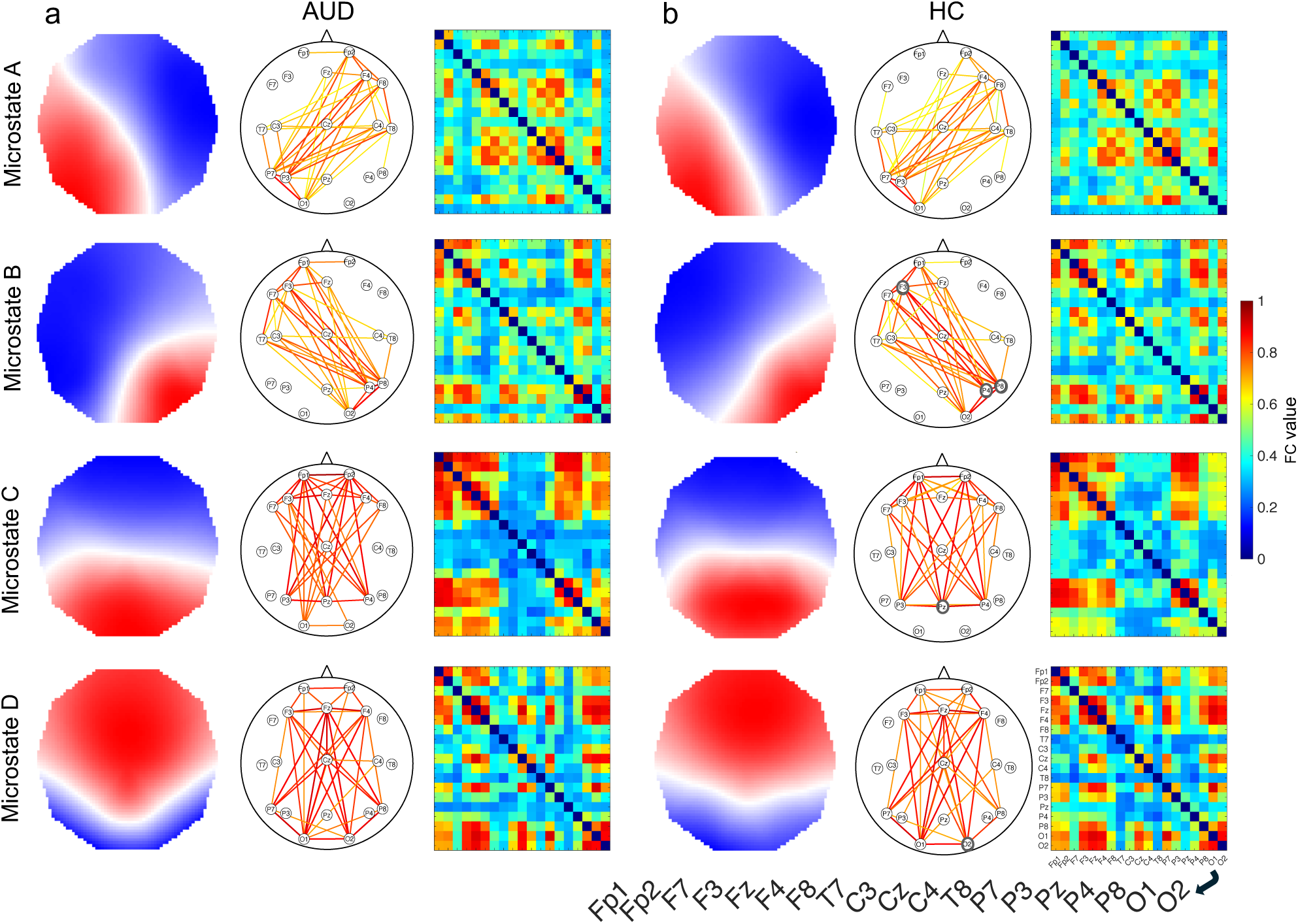
Microstate map topologies corresponded closely with their respective functional connectivity patterns. Four distinct microstates were estimated separately for **(a)** individuals with alcohol use disorder (AUD) and **(b)** healthy controls (HC). For each group, functional connectivity patterns were subsequently computed for each microstate. **(a, b)** The middle plots illustrate the channel pairs showing the top 20% of the strongest functional connectivity within each microstate, with the colors and line thickness indicating connectivity strength. The right plots display all channel pairs.

We assessed temporal dynamics across the four microstates within each group. In the HC group, GEV differed significantly across microstates (*F*(3, 102) = 13.78, *P* < 0.001; **Table 2**). Specifically, microstate D showed significantly higher GEV than microstates A and B (*P* < 0.001), as well as microstate C (*P* = 0.024). In addition, microstate B exhibited higher GEV than microstate A (*P* = 0.021). Time coverage showed a significant main effect of microstate (*F*(3, 102) = 2.98, *P* = 0.034); however, this effect did not remain significant after applying corrections for sphericity (all *P* > 0.05). In addition, no significant differences in mean duration were observed across microstates (*F*(3, 102) = 2.98, *P* = 0.13). These results suggest that, despite differences in GEV, all four microstates occurred with comparable frequency and duration in the HC group.

**Table 2.** Microstate parameters for each state for both participant groups.

|  |  | Microstate A | Microstate B | Microstate C | Microstate D |
| --- | --- | --- | --- | --- | --- |
| GEV | AUD | 0.07 ± 0.04 | 0.09 ± 0.04* | 0.21 ± 0.10* | 0.25 ± 0.13 |
|  | HC | 0.10 ± 0.04 | 0.13 ± 0.04* | 0.14 ± 0.10* | 0.25 ± 0.11 |
| Time coverage (%) | AUD | 19.78 ± 6.46* | 19.61 ± 4.59* | 26.76 ± 8.46 | 33.84 ± 11.68 |
|  | HC | 24.39 ± 6.01* | 24.60 ± 5.04* | 21.80 ± 9.21 | 29.21 ± 9.56 |
| Mean duration (ms) | AUD | 76.04 ± 7.02 | 73.65 ± 6.63 | 85.23 ± 16.20 | 99.69 ± 30.73 |
|  | HC | 80.18 ± 7.27 | 78.47 ± 7.45 | 79.20 ± 26.29 | 89.95 ± 26.10 |

Values are presented as mean ± SD. AUD, Alcohol Use Disorder; HC, Healthy Control; GEV, Global Explained Variance. Parameters that differ significantly ( $P_{FDR} < 0.05$ ) between the AUD and HC groups are marked with asterisks.

In the AUD group, GEV differed significantly across the four microstates (*F*(3, 102) = 21.33, *P* < 0.001), but the pattern differed from that observed in the HC group. Specifically, microstates A and B showed comparable GEV (*P* = 0.29), as did microstates C and D (*P* = 0.75). In addition, microstates C and D each exhibited significantly higher GEV than microstates A and B (all *P* < 0.001). In the AUD group, time coverage and mean duration differed significantly across microstates (time coverage: *F*(3, 102) = 12.69, *P* < 0.001; mean duration: *F*(3, 102) = 9.27, *P* < 0.001). Time coverage of microstate C was comparable to that of microstate D (*P* = 0.22), while both microstates C and D showed higher time coverage than microstates A (C: *P* = 0.028; D: *P* = 0.001) and B (C: *P* = 0.008; D: *P* < 0.001). Similarly, the mean duration of microstate C and D were comparable (*P* = 0.30), whereas both microstates C and D showed longer mean durations than microstates A (C: *P* = 0.046; D: *P* = 0.012) and B (C: *P* = 0.005; D: *P* = 0.003). These results show that, in the AUD group, microstates C and D were more dominant than microstates A and B.

Direct comparisons between the two groups, controlling for participants’ age, revealed significant differences in GEV for microstates B and C, as well as in time coverage for microstates A and B (**Table 2**, **Figure 2**). Specifically, GEV value for microstate B was higher in the HC group than in the AUD group (*t*(59) = 2.86, *P_FDR_* = 0.020), whereas GEV for microstate C was higher in the AUD group than in the HC group (*t*(59) = -2.65, *P_FDR_* = 0.020). In addition, time coverage for microstates A and B was significantly greater in the HC group than in the AUD group (A: *t*(59) = 2.79, *P_FDR_* = 0.013; B: *t*(59) = 3.71, *P_FDR_* = 0.001). These results indicate that the AUD group exhibited dominant and less dominant microstates rather than equally distributed states, suggesting reduced temporal dynamics in their rsEEG.

**Figure 2.**
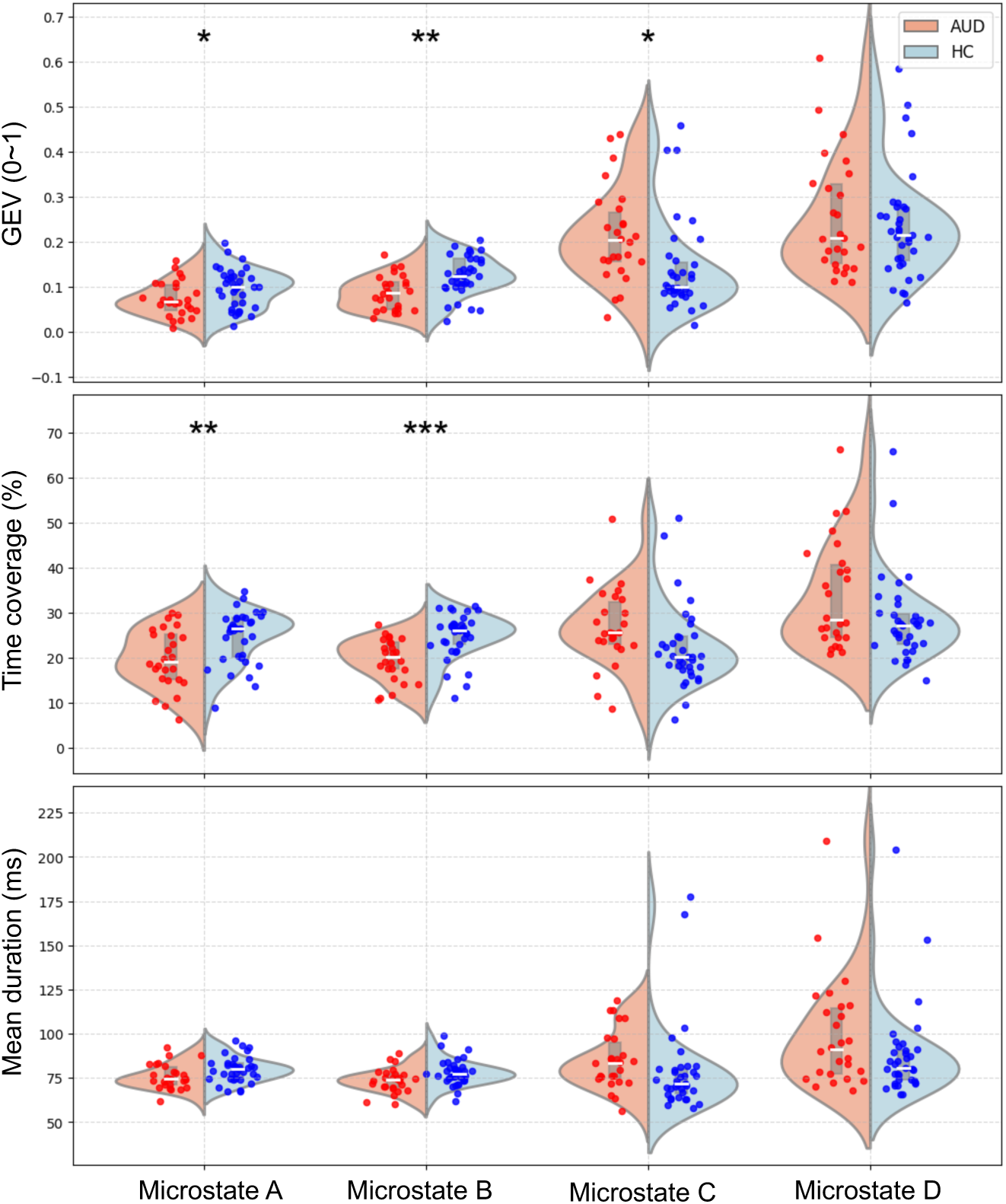
Individuals with alcohol use disorder exhibited significantly reduced time coverage in microstates A and B compared to healthy controls. The global explained variance (GEV) of microstate B was significantly lower in individuals with alcohol use disorder (AUD) than in healthy controls (HC), whereas that of microstate C was significantly higher in individuals with AUD. Consistent with the GEV parameters, time coverage for microstates A and B was significantly shorter in individuals with AUD than in healthy controls. Split violin plots illustrate the distributions of each microstate parameter. Red and blue markers represent individuals with AUD and healthy controls, respectively. Gray box plots indicate the median and the 25th and 75th percentiles. *P_FDR_ < 0.05; **P_FDR_ < 0.01; ***P_FDR_ < 0.001.

### Individuals with AUD showed distinctive microstate-specific functional connectivity patterns compared with healthy controls

Next, we computed functional connectivity between pairs of EEG channels for each microstate and for each group. Each ‘microstate-specific functional connectivity’ pattern matched its corresponding microstate map topology (**Fig. 1**). In both groups, the top 20% of the strongest functional connectivity in microstate A was between channels in the right frontal and left occipital regions; in microstate B, between the left frontal and right occipital regions; and in microstates C and D, between frontal and occipital regions. Subsequently, we conducted PCA-based clustering on functional connectivity across all microstates, which confirmed that microstate-specific functional connectivity patterns were quantitatively separable for both groups (**Supplementary Fig. 1**). These patterns indicate that, as hypothesized, each microstate is associated with a distinct functional connectivity profile.

Comparisons of functional connectivity between the two groups revealed significant functional connectivity differences in microstates C and D (*P_FDR_* < 0.001), but not in microstates A or B (**Fig. 3**). In microstate C, the majority of statistically significant connectivity differences were observed between the frontal and occipital regions, with the AUD group showing hypoconnectivity compared to the HC group. In microstate D, by contrast, the AUD group showed hyperconnectivity compared to the HC group between similar channel pairs.

**Figure 3.**
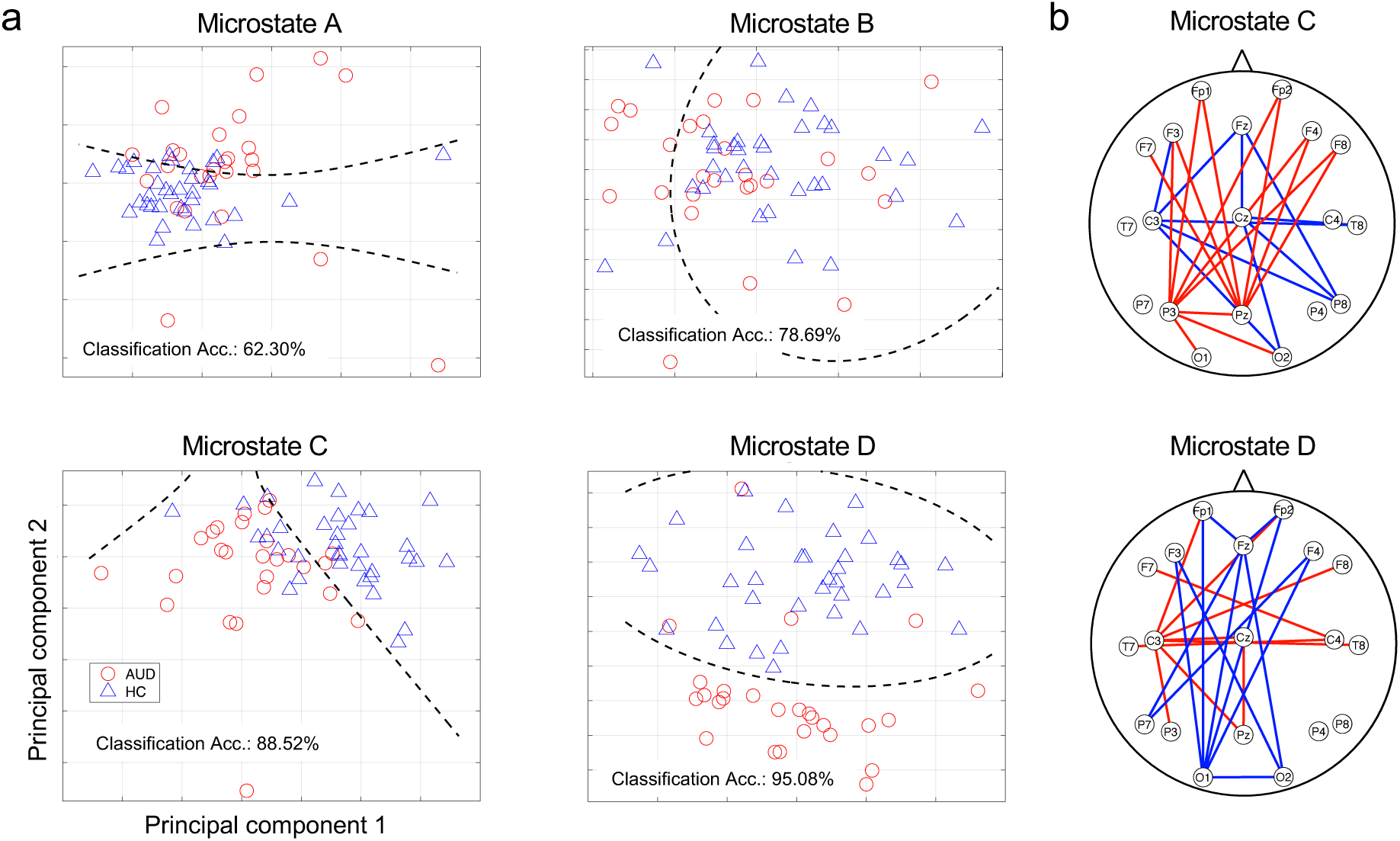
Individuals with alcohol use disorder showed distinctive functional connectivity patterns specific to microstates C and D compared to healthy controls. **(a)** Functional connectivity patterns for each microstate were used to distinguish the two groups using a support vector machine (SVM). Each of the microstate-specific functional connectivity led to classification accuracies higher than chance level. Specifically, patterns from microstates C and D led to higher classification accuracies (88.52% and 95.08%) than those from microstates A and B (62.30% and 78.69%). For illustrative purposes, the functional connectivity values in each group were plotted against their two principal components. The dashed lines indicate the decision boundaries estimated using PCA-based clustering. **(b)** A set of functional connectivity specific to microstates C and D, estimated between frontal and parietal regions, was significantly different between the groups (*P_FDR_* < 0.001). Red lines indicate hypoconnectivity in the alcohol use disorder (AUD) group compared to the healthy control (HC) group, while blue lines indicate hyperconnectivity in the AUD group.

To examine whether EEG measures in the AUD group were associated with psychological status, we computed age-controlled partial correlations between microstate-specific functional connectivity measures that showed significant group differences and individual characteristics, including AUDIT scores, BIS and BAS scores, and PWI scores. In microstate C, functional connectivity between the lateral channel pairs (F8–C3 and F7–C4) showed negative correlations with AUDIT scores (r = -0.40, *P* = 0.049; r = -0.42, *P* = 0.038, respectively; **Fig. 4a**), whereas connectivity between the F4–P7 pair showed a positive correlation (r = 0.42, *P* = 0.038; **Fig. 4a**). However, none of these relationships survived FDR correction. In addition, functional connectivity between the left temporal region and multiple other regions (F8–C3, T7–T8, C3–T8, C3–P3, and C3–Pz) showed significant negative correlations with PWI scores (*Pfdr* < 0.05; **Fig. 4b**). Two other channel pairs, FP2-Fz and F4-O1), showed significant positive correlations between functional connectivity and PWI scores (*Pfdr* = 0.047 and *Pfdr* = 0.029, respectively; **Fig. 4b**). No significant correlations were found with the other psychological variables. In addition, microstate D-specific functional connectivity did not show significant correlations with any of the assessed psychological variables. These significant associations suggest that microstate C-specific functional connectivity is linked to AUD-related symptom severity.

**Figure 4.**
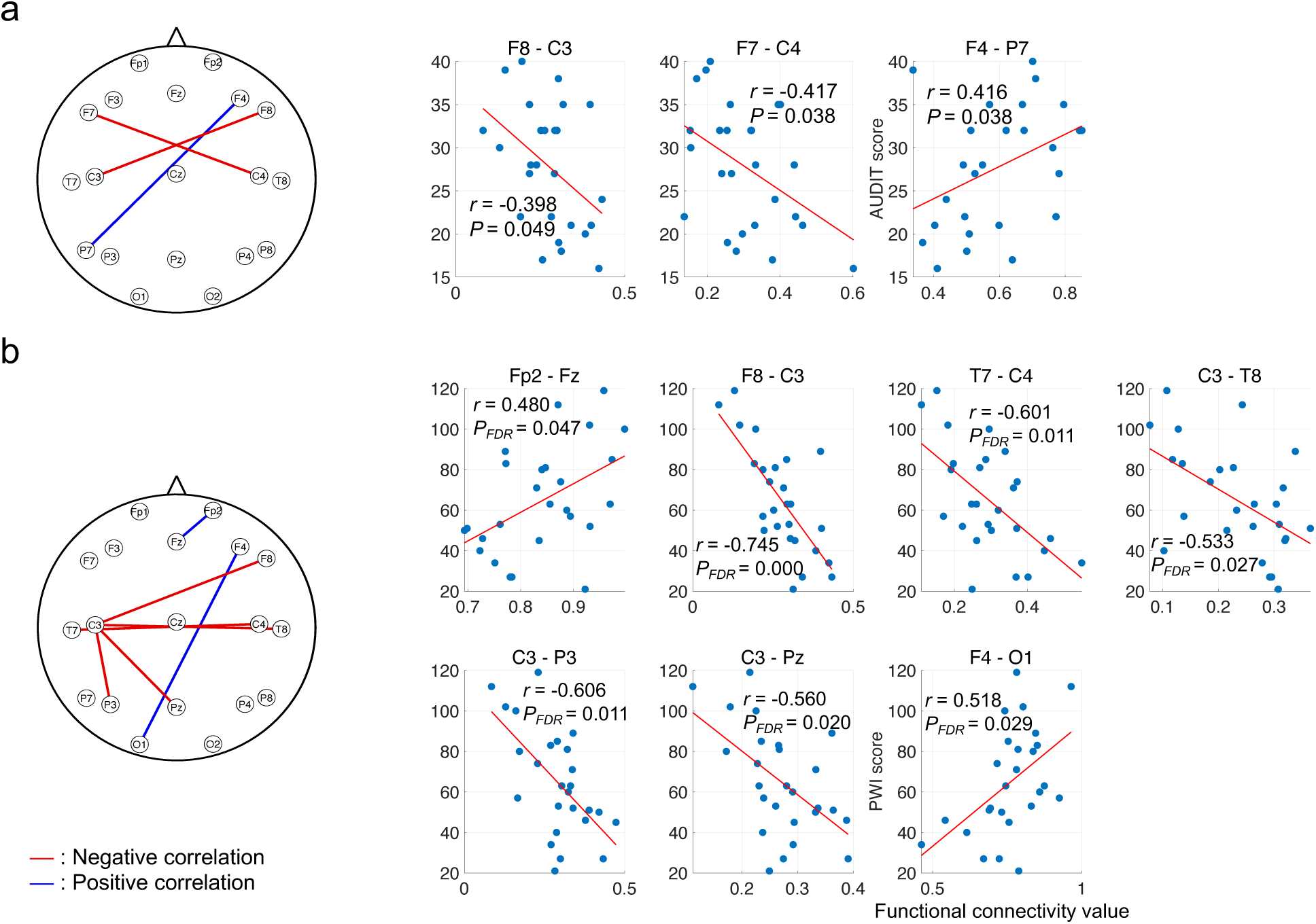
Microstate C-specific functional connectivity in individuals with alcohol use disorder (AUD) is significantly associated with their AUD-related symptoms. (a, **b)** Partial correlations were computed between functional connectivity in individuals with AUD and their AUD-related symptoms, controlling for age. **(a)** Interhemispheric functional connectivity between frontal regions showed significant negative associations with Alcohol Use Disorder Identification Test (AUDIT) scores, whereas the functional connectivity between the right frontal and left parietal channels showed a positive association. **(b)** Functional connectivity between the right frontotemporal or left parietal regions and the left temporal regions showed significant negative associations with Psychosocial Well-being Index (PWI) scores, whereas the functional connectivity between the right frontal and left parietal channels showed a positive association. Each marker represents an individual participant, and the red lines overlaid on the scatter plots illustrate the regression lines.

To directly test our hypothesis that microstate-specific functional connectivity offers a more stable approach for distinguishing HC and AUD groups, we used a support vector machine (SVM) and compared its performance with that based on conventional functional connectivity measures. The SVM based on conventional functional connectivity measures showed very limited discriminative ability (accuracy = 55.74%; **Supplementary Fig. 2**). In contrast, microstate-specific functional connectivity achieved substantially better group separation across all microstates (**Fig. 3a**). Notably, functional connectivity associated with microstates C and D yielded high classification accuracies of 88.52% and 95.08%, respectively. These results indicate that our proposed approach more effectively captures characteristic differences in functional connectivity associated with AUD.

### Putative hub nodes in the AUD network do not show hub-like topological properties

To evaluate the impact of altered connectivity in AUD, we computed network properties of microstate-specific functional connectivity in each group. Specifically, we calculated network strength for all channel nodes and identified nodes exceeding one standard deviation above the mean as hub candidate nodes (**Fig. 5**). By examining consistencies over four additional network properties (EC, C, BC, and CC; see **Methods**), we identified channel nodes that showed hub-like properties. In the HC group, hub-like nodes were identified from microstates B (F3, P4, and P8), C (Pz), and D (O2). However, in the AUD group, BC and CC values of the candidate nodes did not meet the criteria for a hub identification, and therefore no hub nodes were identified in the AUD group (**Fig. 5**). These results—showing that the candidate nodes had low BC and high CC—indicate that the functional connectivity network in AUD is more locally connected, and suggest potentially impaired global communication.

**Figure 5.**
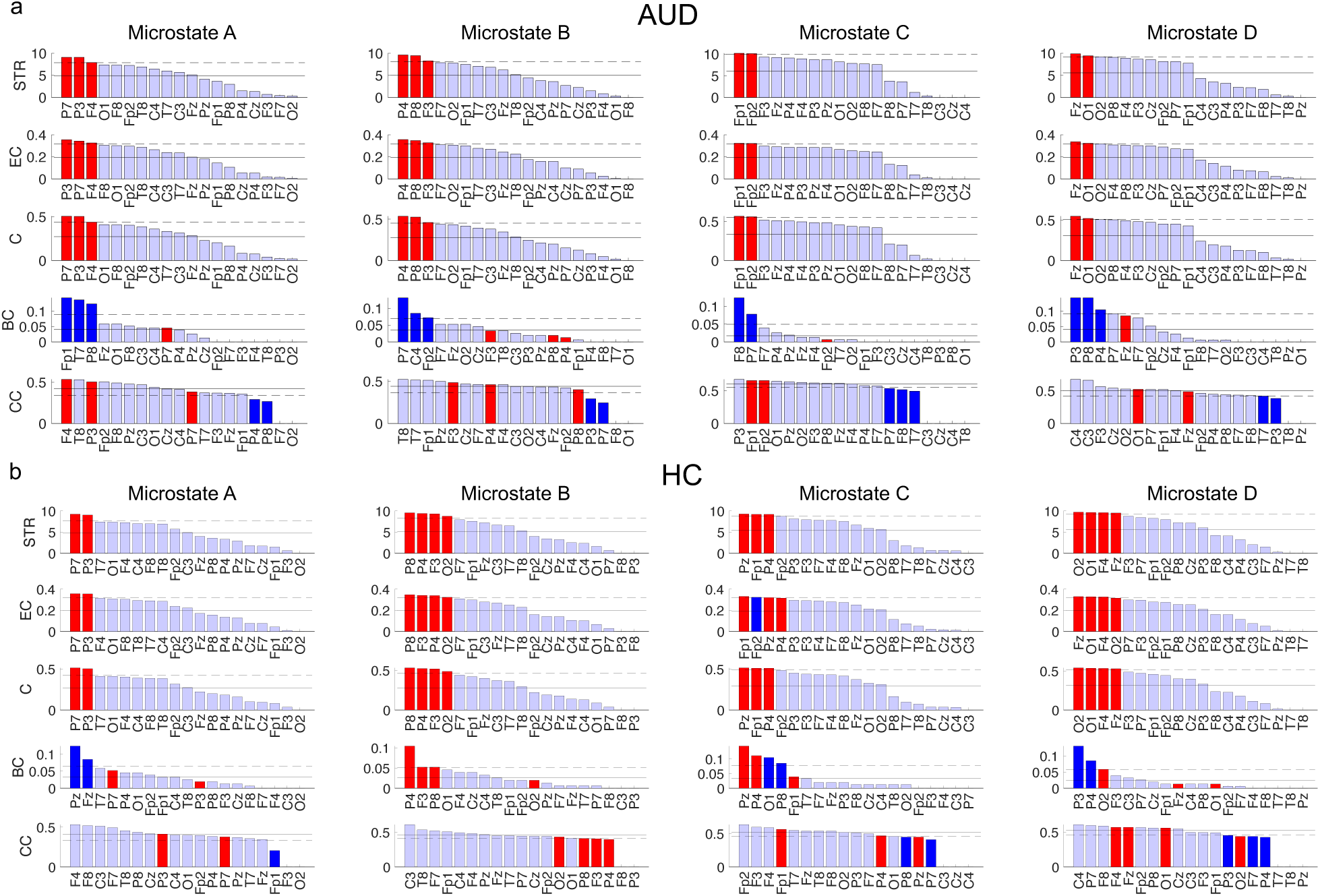
rsEEG network nodes in individuals with AUD showed less stable hub-like properties compared with healthy controls. **(a, b)** EEG channels were defined as network nodes, and graph-theoretical features were computed to characterize each group’s neural network properties. For each group and each microstate network, nodes with network strength (STR) exceeding one standard deviation above the mean were identified as hub-like nodes (red bars). Nodes are displayed in descending order according to their eigenvector centrality (EC), closeness centrality (C), betweenness centrality (BC), and clustering coefficient (CC), respectively. Nodes that did not meet the hub criterion based on STR but exceeded the one-standard-deviation threshold (above thresholds for EC, C, and BC, while below the threshold for CC) for each respective feature are highlighted in blue. Solid and dotted lines indicate the mean and one-standard-deviation threshold, respectively. The identified nodes showed stable hub-like properties across EC and C in both groups. **(b)** In healthy controls, particularly for microstates B, C, and D, these nodes also exhibited stable properties across BC and CC, **(a)** whereas the identified nodes in the AUD did not.

In addition, we evaluated the network efficiency of each microstate-specific connectivity network using the small-worldness metric (**Fig. 6**). Compared to the HC group, the AUD group showed significantly lower small-worldness in the microstate C-specific network (HC: 1.03 ± 0.04; AUD: 1.00 ± 0.03; *t*(59) = -3.71, *P_FDR_* = 0.001). Small-worldness did not differ between groups for the other microstate-specific networks (A, HC: 1.04 ± 0.04; AUD: 1.06 ± 0.06, *P* = 0.27; B, HC: 1.03 ± 0.04, AUD: 1.06 ± 0.06, *P* = 0.078; D, HC: 1.04 ± 0.03; AUD: 1.03 ± 0.03, *P* = 0.10). This result indicates that individuals with AUD have a less efficient network structure, specifically in microstate C.

**Figure 6.**
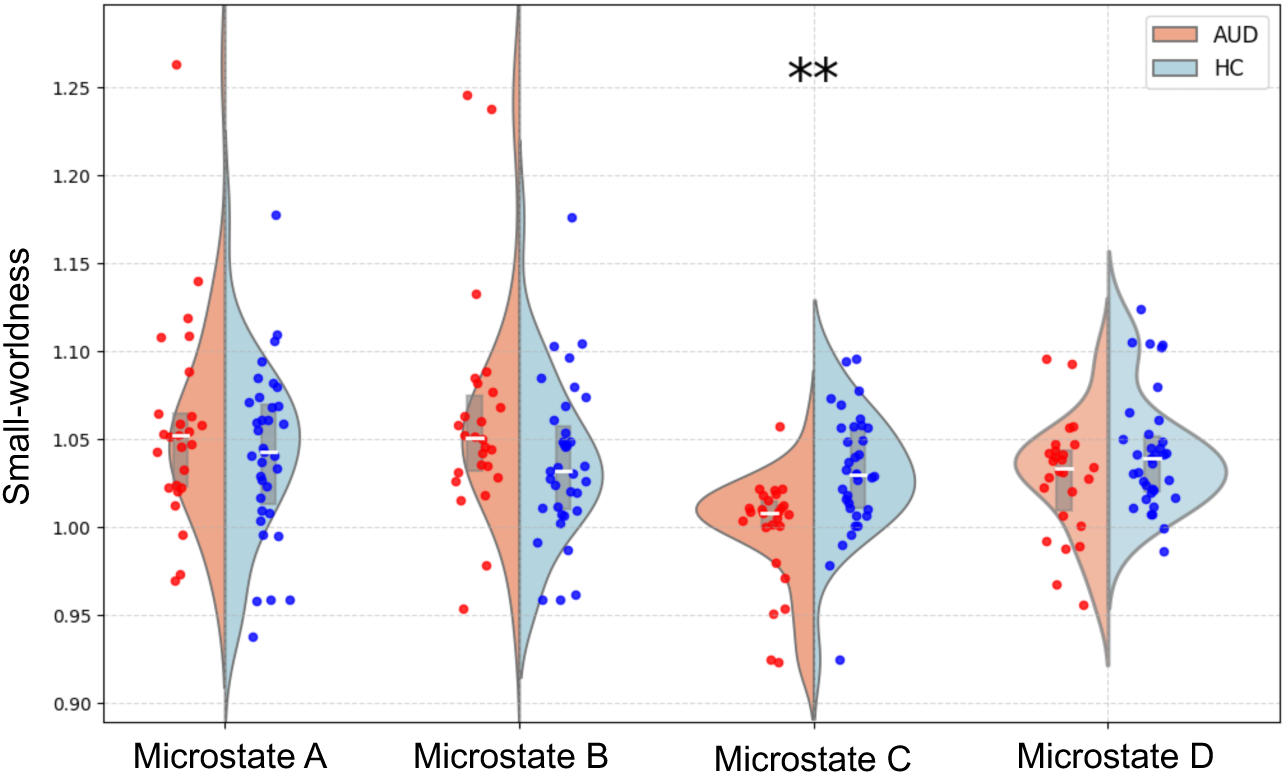
The microstate C-specific functional connectivity network in individuals with AUD showed significantly lower small-worldness than that of healthy controls. To examine the efficiency of each network, small-worldness was computed for each group and for each microstate-specific functional connectivity network. Only for microstate C, the small-worldness of the network was significantly lower in individuals with AUD compared with healthy controls (*P_FDR_* = 0.001). Split violin plots illustrate the data distributions, and each marker represents an individual participant. Gray box plots denote the median along with the 25th and 75th percentiles. ** *P_FDR_* < 0.01.

## Discussion

In the current study, we investigated functional connectivity in individuals with AUD while accounting for transient state changes in their resting-state EEG. Although the microstate map topologies were comparable between groups, the temporal dynamics across the four microstates showed significant differences. Moreover, compared to functional connectivity computed using a conventional approach, functional connectivity specific to microstates C and D demonstrated superior discriminative power in distinguishing the AUD group from the HC group. These results are in line with our hypothesis that failing to account for dynamic state changes contributes to inconsistencies in functional connectivity measures, and they highlight the potential utility of microstate-specific functional connectivity for investigating AUD.

The microstate time series has been known to vary across neuropsychiatric disorders (Kikuchi et al., 2011; Kim et al., 2024; Lehmann et al., 2005). In contrast to the classical assumption of stationarity in EEG, distinct topologies of electric potentials can be defined, whose transient occurrences and transitions capture global representations of the brain’s dynamic states and their abnormalities (for review, see Khanna et al., 2015). Our data showed that the AUD and HC groups did not differ in the topological maps of the identified microstates, but rather differed in their occurrence patterns. Increased dominance of microstates C and D—transient states associated with the default mode network (DMN) and executive control network (ECN), respectively, rather than low-level sensory networks (for review, see Tarailis et al., 2024)—over other microstates indicates diminished network dynamics and reduced network efficiency in AUD. This pattern may reflect persistent self-referential processing accompanied by compensatory recruitment of executive control systems, a profile commonly reported in addictive disorders (Goldstein and Volkow, 2011; Zilverstand et al., 2018). Note that our data were derived from rsEEG, and thus interpretations regarding the functional impact of dominant microstates are necessarily speculative and based on inferences from prior literature.

Functional connectivity analysis combined with microstate segmentation revealed distinct connectivity alterations in individuals with AUD between microstates C and D. Specifically, functional connectivity during microstate C was significantly reduced in the AUD group, consistent with the notion of a disconnection syndrome (Dupuy and Chanraud, 2016). In contrast, connectivity patterns during microstate D were mixed, showing increased connectivity in some channel pairs and decreased connectivity in others. Consistent with previous literature across different neural measure modalities emphasizing the importance of dynamic connectivity analyses (Baker et al., 2014; Hutchison et al., 2013; Michel and Koenig, 2018), our microstate-specific connectivity results were more informative in distinguishing individuals with AUD from healthy controls. These findings suggest that inconsistencies reported in previous AUD studies may, in part, reflect the lack of separation among distinct microstates. It is worth noting that, in addition to local connectivity alterations (i.e., functional connectivity between channels), the global functional network in the AUD group was less efficient, as evidenced by the absence of stable hub-like properties. While our results point to a potential association between alcohol-related impairments in local, microstate-specific connectivity and reduced global network efficiency, the mechanistic pathways linking these effects are not yet understood, underscoring an important direction for future investigation.

The current study has the following limitations. First, the demographic characteristics of the AUD and HC groups were not fully matched; in particular, the AUD group was significantly older than the HC group. Although age was controlled for in all between-group statistical analyses, future studies with better-matched samples are needed to confirm the replicability of these findings. Second, the sample sizes of our AUD and HC groups were relatively small to modest. Given the high dimensionality of EEG-related measures, all statistical tests in our study were corrected for multiple comparisons, ensuring that only statistically robust findings were reported. Still, future studies with larger sample sizes would be important for examining individual differences and for evaluating microstate-specific functional connectivity measures as potential neural markers of alcohol-related characteristics. Third, interpretations of the observed between-group differences in microstate-specific functional connectivity remain speculative, as they are based on inferences drawn from prior literature. To address this limitation, future studies could adopt longitudinal designs or causal neuromodulation approaches to track changes in individuals’ alcohol dependence alongside corresponding alterations in EEG features.

Nevertheless, our findings suggest that EEG microstate-specific functional connectivity provides informative neural markers for characterizing the neural impact of alcohol use in young adults. The proposed approach to examining dynamic networks and their graph-theoretical characteristics delineated neural patterns that converge with prior empirical reports of brain’s structural and cognitive deficits in AUD (Rao and Topiwala, 2020; Shokri-Kojori et al., 2017), helping to distinguish AUD-related alterations from other mixed patterns. In sum, these findings contribute to understanding how uncontrolled alcohol use is related to both local and global brain network organization, and they highlight the potential utility of transient EEG properties as a tool for monitoring the severity of alcohol addiction.

## Acknowledgements

This work was supported in part by the National Research Foundation of Korea (NRF) (RS-2024-00420674 to D.C. and J.-S.C.; RS-2025-02263045 to D.C.).

## Author contributions

Hanjin Park: Data curation, Formal analysis, Visualization, Writing – original draft, Writing – review and editing;

So Young Yoo: Data curation, Investigation, Writing – review and editing;

Hyunho Lee: Data curation, Investigation, Writing – review and editing;

Mnkyung Park: Data curation, Investigation, Writing – review and editing;

Jung-Seok Choi: Funding acquisition, Supervision, Writing – review and editing;

Dongil Chung: Conceptualization, Funding acquisition, Supervision, Writing – original draft, Writing – review and editing.

