## Supplementary figures for "Microstate-specific functional connectivity in alcohol use disorder using resting-state EEG"

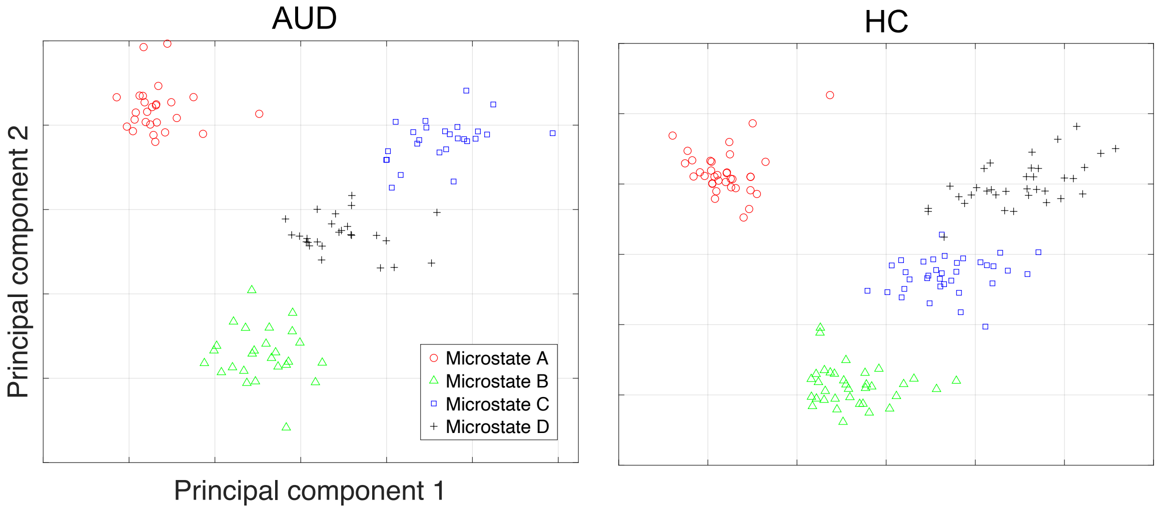


**Supplementary Figure 1. Results of PCA-based clustering of microstate-specific functional connectivity patterns for the AUD and HC groups.** In both groups, connectivity patterns corresponding to each microstate were clearly distinguishable. Although the connectivity patterns of microstates C and D were more similar to each other—and thus located closer in the PCA space—than to the other microstates, they remained clearly separable. In contrast, microstates A and B showed the greatest separation, consistent with their map topologies.

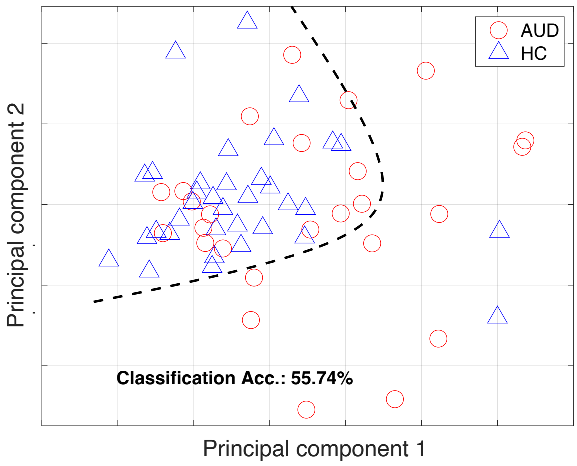


**Supplementary Figure 2. Classification results for the AUD and HC groups using conventional functional connectivity estimation.** PCA-based clustering of conventional functional connectivity patterns and the corresponding classification accuracy are shown. The dashed line denotes the decision boundary. Substantial overlap between data points from the AUD and HC groups resulted in classification accuracy at the chance level. These findings indicate that conventional functional connectivity estimation has limited sensitivity for detecting AUD-related alterations in functional connectivity.
